# Genomic surveillance and phylodynamic analyses reveal emergence of novel mutation and co-mutation patterns within SARS-CoV2 variants prevalent in India

**DOI:** 10.1101/2021.03.25.436930

**Authors:** Nupur Biswas, Priyanka Mallick, Sujay Krishna Maity, Debaleena Bhowmik, Arpita Ghosh Mitra, Soumen Saha, Aviral Roy, Partha Chakrabarti, Sandip Paul, Saikat Chakrabarti

## Abstract

Emergence of distinct viral clades has been observed in SARS-CoV2 variants across the world and India. Identification of the genomic diversity and the phylodynamic profiles of the prevalent strains of the country are critical to understand the evolution and spread of the variants. We performed whole-genome sequencing of 54 SARS-CoV2 strains collected from COVID-19 patients in Kolkata, West Bengal during August to October 2020. Phylogeographic and phylodynamic analyses were performed using these 54 and other sequences from India and abroad available in GISAID database. Spatio-temporal evolutionary dynamics of the pathogen across various regions and states of India over three different time periods in the year 2020 were analyzed. We estimated the clade dynamics of the Indian strains and compared the clade specific mutations and the co-mutation patterns across states and union territories of India over the time course. We observed that GR, GH and G (GISAID) or 20B and 20A (Nextstrain) clades were the prevalent clades in India during middle and later half of the year 2020. However, frequent mutations and co-mutations observed within the major clades across time periods do not show much overlap, indicating emergence of newer mutations in the viral population prevailing in the country. Further, we explored the possible association of specific mutations and co-mutations with the infection outcomes manifested within the Indian patients.

## Introduction

The corona virus disease or COVID-19 pandemic caused by the SARS-CoV2 virus has created an unprecedented health and financial crisis throughout the world (1, 2). Since the emergence of the outbreak in the Chinese city of Wuhan in late 2019, the novel corona virus disease has spread widely and has caused millions of infections and thousands of death throughout the year 2020 and active infections are still continuing at an alarming rate in parts of the globe, especially in Americas and Europe. As of March 2021, the total number of infections reported in India surpasses 11 million while active infections are still more than 180,000 causing overall death toll more than 158,000 (3). Similar to global efforts to combat the deadly disease, various measures have been taken by the Indian clinical and biomedical research community including vaccine development (4), clinical trials with repurposed drugs (5), convalescent plasma therapy (6), and genetic surveillance via genome sequencing of the viral strains extracted from the infected individuals (7–15).

Several thousands of whole-genome sequences of the SARS-CoV2 from various parts of the country have been sequenced and subsequently deposited in global databases such as GISAID (16). Multiple works from India have highlighted the genomic diversity and the phylogenetic profiles of the prevalent strains in the country (7, 17). However, more than 75% of the SARS-CoV2 sequences (2521 out of 3277 complete genomes from India) were deposited in the latter half of the year 2020. It is understandable for the lower number (146) of the deposited sequences during December 2019 to March 2020 as the active cases only started to appear in India in March 2020 only. In order to enrich the viral genome sequence data for the latter half of the year, we decided to sequence 54 SARS-CoV2 sequences collected from the state of West Bengal during August 2020 to December 2020 making it only the third Indian state apart from Telangana and Maharashtra to deposit more than fifty sequences in that time period. With these sequences we decided to analyze and understand the spatio-temporal evolutionary dynamics of the pathogen across various states and union territories (UT) of India in the year 2020. We estimated the clade dynamics of the Indian strains and compared the clade specific mutations, speculated their positive selection and calculated the co-mutations patterns across states and UTs of India. Further, we explored the possible association of specific mutations/co-occurred mutations with the infection outcome manifested within the patients.

We found that for Indian sequences GR, GH and G (GISAID) or 20B and 20A (Nextstrain) (18) clades were primarily the major prevalent clades in middle and later half of the year 2020. However, frequent mutations observed within each of the major clades do not show much overlap, especially for the last half of the year 2020, indicating the emergence of new mutations in the viral population prevailing in the country. Interestingly, only 10% of the mutations within the GISAID clades across various Indian states are found to be common. Co-mutations or co-occurrence of mutations within a specific viral strain were investigated and frequent co-mutation patterns for different Indian states were identified. Finally, associations between specific mutation and co-mutation pattern with respect to patient status (deceased, symptomatic, asymptomatic, etc) have been explored.

## Materials and Methods

### Sample collection

Ethical clearances were taken from the Institutional (CSIR-Indian Indian Institute of Chemical Biology) and hospital (MEDICA Superspecialty Hospital) ethical committees for the present study. Nasopharyngeal/ oropharyngeal swabs of COVID-19 patients were collected from August to October 2020. Samples were anonymized by removing patient identifiers except gender, age, and collection date. SARS-CoV-2 nucleic acids were isolated using the MagMax Viral Pathogen Isolation kit from Themo Fisher in KingFisher Flex automated extractor. standard protocol. The RT-PCR assay was performed using SD Biosensor COVID 19 kit based on taqman probe chemistry for the detection of SARS-CoV2 RDRP gene and E gene using reverse transcription in Rotor Gene Q 5 plex HRM system. Samples with Ct values <= 25 were considered for sequencing.

### Viral whole genome sequencing

Virus genomes were sequenced using ARTIC COVID-19 multiplex PCR primers version 3 by via combination of nanopore sequencing based on MinION sequencer, Oxford Nanopore Technology (ONT) and Illumina HiSeqX (19). To generate PCR amplicons for nanopore sequencing Native Barcode Expansion 1-12, protocols (Kits EXP-NBD104) (ONT) were used. RNA extracted from the clinical specimens were converted to cDNA using reverse transcriptase enzyme (Super Script IV First Strand Synthesis Kit, Thermo Fisher Scientific, Waltham, MA) and then purified by using AMPure XP beads. Purified cDNA was then amplified by each of the two ARTIC v3 primer pools which tile the SARS-CoV2 genome. The amplified product was further subjected to end-repair, and barcoded by Native Barcode Expansion kits (1–12) and purified by 0.4X of concentration of AmpureXP. All samples were then pooled together and ligated with sequencing adapters, purified samples were finally quantified using Qubit 4.0 Fluorometer (Invitrogen), followed by loading of 15ng of pooled barcoded material and sequencing on MinION flow cells.

For Illumina sequencing, samples were sequenced through 2×150bp paired end Illumina’s HiSeqX sequencing system following standard protocol. QIAseq SARS-CoV-2 Primer Panel (Qiagen, cat. no. 333895) and QIAseq FX DNA Library Kit were used in order to prepare amplicon libraries for viral genome sequencing. Prepared libraries were then pooled and sequenced using Illumina HiSeqX instrument to generate 150bp paired end reads.

### Genome assembly and sequence submission in GISAID

Two different methods were implemented for the assembly of long and short reads as obtained from ONT and Illumina sequencing platforms, respectively. All reads from both the platforms were checked for their quality using FastQC (20). In case of long reads, reference based assembly was performed using the first SARS-CoV2 strain identified from China, Wuhan (NCBI Accession Number NC-045512.2 (21) which is identical to the GISAID reference sequence EPI_ISL_402124 (16)) as reference, and Minimap2 (22) with ONT specific parameters. In case of short reads, they were first filtered using kneaddata (23) and then assembled by SPAdes (24) using default parameters. Pilon (25) was used for polishing and generation of final consensus sequences. The assembled SARS-CoV2 genome sequences were checked for frameshifts using Genome Detective online tool (26) and their depths were calculated using Mosdepth (27). All the 54 genome sequences with their associated metadata were uploaded to GISAID database (Table S1).

### Sequence and mutation data collection

We have accessed the protein sequences of SARS-CoV2 virus collected from different continents from the EpiCoV database of GISAID (16). The database was searched on 1^st^ January 2021 up to sample collection date 31^st^ December 2020 using the primary key-words ‘hCoV-19’ and ‘Human’. Only complete and high coverage sequences were considered. Sequences with genomes >29,000 bp were considered complete.

Sequences with <1% Ns (undefined bases) were considered as high coverage sequences. Sequences collected from India were analyzed separately. Sequences from different states of India were also accessed and analyzed separately. Additional metadata for the sequences which include location of sample collection, age and sex of the patients, clade, lineage, patient status were also downloaded.

We found only 23 sequences collected in 2019, all from Wuhan, China from where the disease spread. To analyze the evolution of viral clades with time, we divided all sequences in three terms, depending on the date of collection of sample. ‘Term1’ includes sequences collected till March 2020. ‘Term2’ defines April 2020 to July 2020 and ‘Term3’ includes sequences collected from August 2020 to December 2020. Table 1 shows the number of sequences collected from different continents and India in different terms.

**Table 1:** Number of sequences deposited in the year 2020.

| **Continent/Country/Lab** | **December 2019**<br>**– March 2020** | **April 2020 –**<br>**July 2020** | **August 2020 –**<br>**December 2020** | **Total** |
| --- | --- | --- | --- | --- |
| North America | 10877 | 21895 | 8583 | 41355 |
| South America | 433 | 1076 | 308 | 1817 |
| Europe | 16198 | 30827 | 86563 | 133588 |
| Africa | 180 | 1670 | 1019 | 2869 |
| Asia* | 3178 | 9189 | 1249 | 13616 |
| Oceania | 2001 | 6604 | 4184 | 12789 |
| India | 146 | 2521 | 610 | 3277 |
| IICB<sup>#</sup> | 0 | 0 | 54 | 54 |

**\*Excluding India**
**# Indian Institute of Chemical Biology submitted 54 out 55 total sequences deposited in GISAID during August 2020 – December 2020.**

For identifying the mutations, GISAID reference sequence EPI_ISL_402124, collected from human sample in Wuhan, China, in December 2019, was considered as reference (28).

### Alignment and mutation frequency analysis

To extract the unique representative sequences and exclude redundant sequences CD-HIT (29) server was used. The number of CD-HIT runs was kept as 1 with sequence identity cut-off 1.0 (100% identity). It provided clusters of sequences which are 100% identical. The cluster representative sequences along with the reference sequence were aligned using Kalign protein sequence alignment tool (30). Python (version 3.4) codes were used for extracting mutations and further analysis. Mutation was considered as frequent when frequency was calculated with at least 50 sequences (N) and the frequency was ≥2.5% for N ≥200 and mutation count was at least 5 when N < 200.

### Mutational and Co-mutational analysis

For mutational analysis within India, we have chosen states and UTs, which had at least 50 sequences in a given term. For Term2, we found 8 states and UTs. Among them, only 3 states had more than 50 sequences in Term3. Hence, temporal analysis was done for those 3 states only. We also analyzed how different mutations co-occurred in different states and terms. Network was constructed for each state and UT showing co-occurrence between frequent mutations of that state/UT. Cytoscape.js (31) was used to construct the network.

### Patient status association analysis

To correlate disease severity with mutation and co-mutation pattern, available metadata was analyzed. Based on the available ‘patient status’, patients were classified broadly in four categories, ‘deceased’, ‘symptomatic’, ‘mild’, and ‘asymptomatic’. Patient status was associated with frequent mutations, co-occurring mutations, and clades. Fisher’s Exact test was performed using the following contingency table (32) for deceased samples,

|  | **Mutated** | **Not-mutated** | **Total** |
| --- | --- | --- | --- |
| **Not deceased** | a | b | a+b |
| **Deceased** | c | d | c+d |
| **Total** | a+c | b+d | a+b+c+d = N |

where *N* is the total number of sequences. Similar tables were used for other types of patients’ categories (‘symptomatic’, ‘mild’, and ‘asymptomatic’). The probability of obtaining a given set of result, *p-value*, is provided by a hypergeometric distribution,

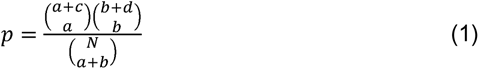

where 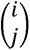 denotes binomial coefficient of any given variable *i* and *j*.

## Results

### Demographic and phylogenic distribution

54 SARS-CoV2 samples taken from patients residing in Kolkata and West Bengal were sequenced using whole genome sequencing approach. The age of the patients ranges from 6 to 88 years with a maximum peak (24.07%) in the range of 51-60 years (Figure S1A) with an overall male to female ratio of 1.7 against the national ratio of 1.99. For comparison we have also plotted the age distribution of the Indian patients for which the extracted SARS-CoV2 sequences were deposited in GISAID database (Figure S1A). The quality of the sequencing data represented in the form of depth shows a mean depth of approximately 26035X and 455X for short reads and long reads respectively (Figure S1B) while the coverage for all the assembled genomes was above 99%. The GISAID clade distribution of the 54 sequences is slightly different from the national distribution with majority of O clade representation and same is true for Nextstrain clades with prevalence of 20A clade (Figure S1C-D).

### Dynamics of clade distribution

In order to explore the evolutionary and mutational dynamics represented in terms of phylogenetic clades we compared the clade distribution of all the complete and high-quality SARS-CoV2 sequences (3277) deposited in GISAID from India until December 31, 2020. The sequences were categorized into three time points (‘Term’) based on the date of the collection/submission of the sequences. Figure 1 shows the distribution of different GISAID and Nextstrain clades in three time points. It is evident that GR clade is prevalent in Term2 and Term3 followed by GH and G clades (33). Similarly, Nextstrain clades 20B and 20A are found to be more prevalent in the latter half of the year 2020. To compare the clade dynamics with respect to SARS-CoV2 sequences from elsewhere, we have performed the similar analysis with sequences deposited from North America, South America, Europe, Africa, Oceania and Asia (without Indian sequences) (Figure S2). Interestingly, except North America and Europe, all the other continents show prevalence of GR clade in the second and third term of the year. North America shows a consistent prevalence of GH clade throughout the year where a massive increase in the numbers of GV clade sequences was observed in Europe during the latter part of the 2020. However, it is interesting to investigate whether sequences belonging to the major clades observed in India through various time ‘Terms’ harbor similar mutations or not. Hence we compared the non-clade defining and ‘frequent’ (frequency >=2.5% or >5 when N <= 200) mutations observed within the same clade across the time terms (Figure S3). Interestingly, we observed that a significant fraction of the mutations turned out be unique in ‘Term2’ and ‘Term3’ indicating accumulation of novel mutations.

**Figure 1:**
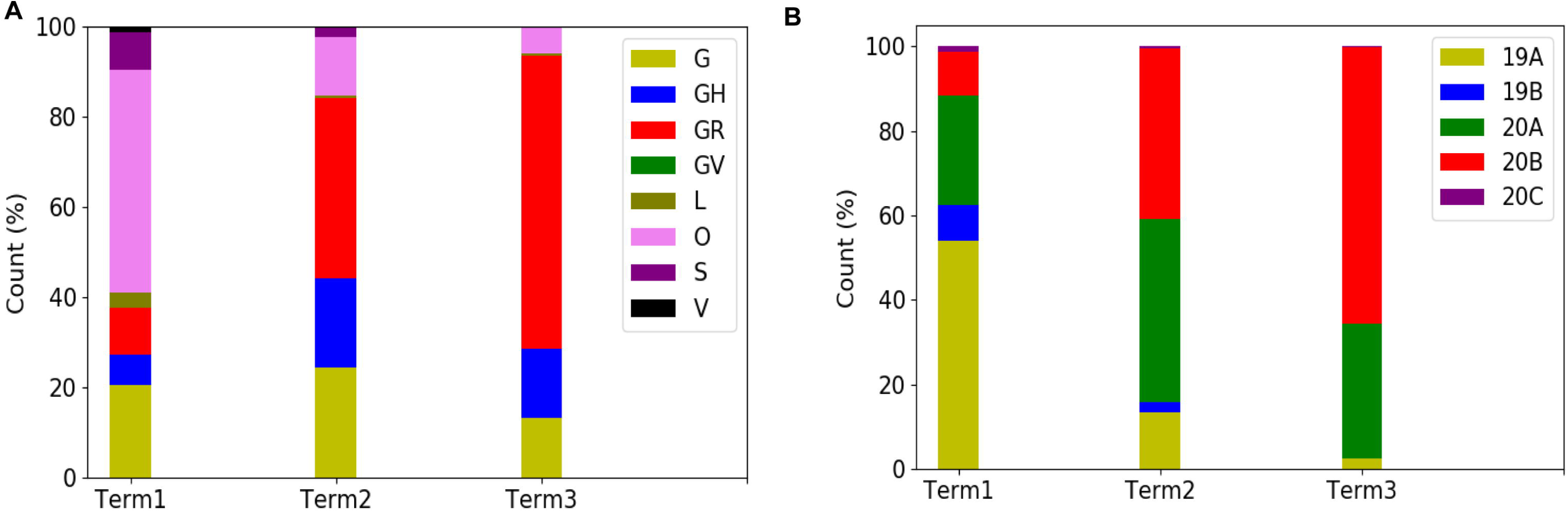
Distribution and dynamics of SARS-CoV2 clades in India for three different time spans in the year 2020. Distribution of GISAID (panel A) and Nextstrain (panel B) clades across India for three different times spans, ‘Term1’ (December 2019 to March 2020), ‘Term2’ (April 2020 to July 2020) and ‘Term3’ (August 2020 to December 2020), respectively.

### Dynamics of clade variation across Indian states and union territories

Curious to see the previous observation, we wanted to investigate whether the mutational variability within clades is more specific to certain geographical location delineated by the states and UTs of India. ‘Term1’ contains only 146 sequences from India, hence, we compared state specific mutations taking SARS-CoV2 sequences from only ‘Term2’ and Term3’, respectively. Only seven states and one union territory, Karnataka (KA), Telangana (TG), Maharashtra (MH), Gujarat (GJ), Delhi (DL), Uttarakhand (UT), West Bengal (WB) and Odisha (OR) contained more than 50 deposited good quality sequences in GISAID. Figure 2A shows fraction of various GISAID clade sequences within these states and union territory. MH, GJ and WB are the top three states representing G clade sequences in ‘Term2’ whereas GJ produces a significantly higher number of GH clade sequences compared to other states. MH and TG possess maximum number of GR sequences followed by KA and OR. Interestingly, DL and TG provides maximum number of O clade sequences which are generally less clearly defined (34). Distribution of clade S is underrepresented in these eight states and union territories. Comparison of the frequent mutations across states for the same clade also revealed much more specific mutation than common ones (Figure 2) where overall only 10% of the mutations within the GISAID clades across various Indian states are found to be common (Table 2). In ‘Term3’ only from three states (MH, TG and WB) more than fifty sequences were deposited and similar to observation from ‘Term2’, very few overlap of mutations were found to be common across these three states (Figure S4 and Table 2).

**Figure 2:**
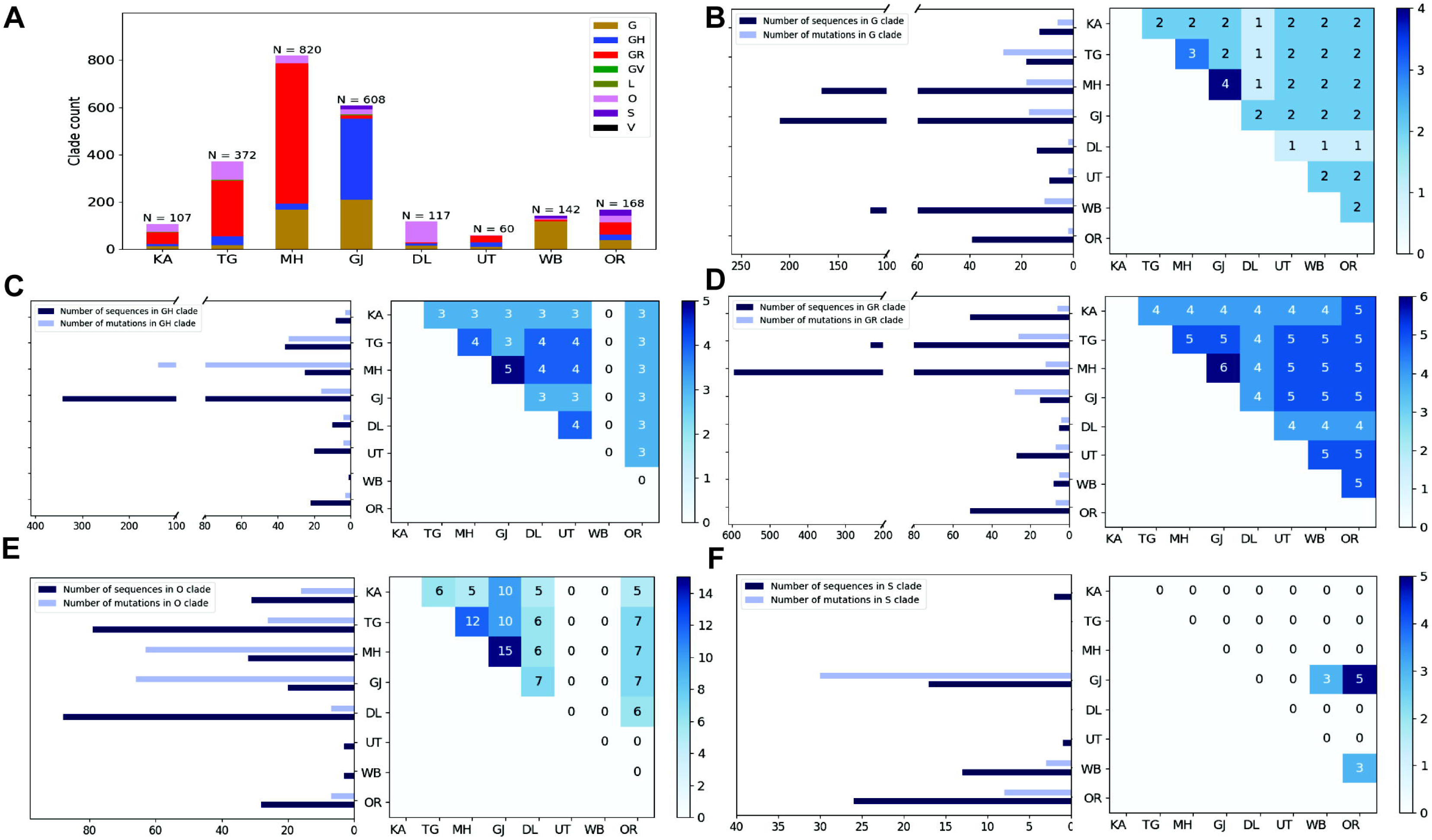
Distribution of GISAID clades within Indian states and union territories and comparison of frequent mutations across them during Term2 (April 2020 to July 2020). Panel A shows the distribution of GISAID clades in seven Indian states and one union territory (UT) that deposited more than 50 SARS-CoV2 sequences during April 2020 to July 2020. Panels B-F plot the number of sequences and frequent mutations of the seven states and one UT and show the number of common mutations among them for G, GR, GH, O, and S clades, respectively. KA: Karnataka, TG: Telangana, MH: Maharashtra, GJ: Gujarat, DL: Delhi, UT: Uttarakhand, WB: West Bengal, OR: Odisha.

**Table 2:** Specific and common mutations across different states for different clades

| **Clades** | **Unique specific mutations in Term2** | **Unique common mutations in Term2** | **Unique specific mutations in Term3** | **Unique common mutations in Term3** |
| --- | --- | --- | --- | --- |
| G | 68 | 6 | 0 | 2 |
| GH | 179 | 6 | 5 | 5 |
| GR | 59 | 7 | 21 | 6 |
| GV | 0 | 0 | 0 | 0 |
| L | 17 | 5 | 0 | 0 |
| O | 133 | 24 | 0 | 0 |
| S | 30 | 5 | 0 | 0 |
| V | 1 | 0 | 0 | 0 |

### Variation in the co-occurrence of mutations (co-mutations) across Indian states and union territories

Combined impact of co-occurrence of mutations within a specific viral strain could be crucial to elicit infectivity and sustenance of viral load within the host. Hence, co-mutation patterns within SARS-CoV2 variants were investigated and specific/common co-mutations for different Indian states were identified irrespective of clade. Figure 3A-3H shows network representation of co-mutations patterns amongst the most frequent mutations where each node represents a mutation site and the edge denotes co-occurrence between a pair of mutant sites. Edge thickness is proportional to the number of co-occurrence while node size is to the frequency of mutation. Any two nodes (sites) were considered to be co-mutated if the co-mutation pair is represented in >=2.5% of the population size or >5 when the total number of sequences in the particular category is less than 200. Figure 3I-3J show the frequency distribution of the number of co-mutating sites for viral strains from different states and union territories. It is evident that for most of these states and UT a large fraction of the sequences harbor five or more co-mutations. Although the mutations include the clade defining mutations [e.g., G: Spike(D614G), GH: Spike(D614G) and NS3(Q57H), GR: Spike(D614G) and N(G204R), GV: Spike(D614G) and Spike(A222V), S: NS8(L84S)] a large number non-clade defining mutations are found to co-occur within SARS-CoV2 strains from these states and UT during the ‘Term2’. In ‘MH’ more than 50% sequences had 7 co-mutations. ‘KA’ also has similar trend. However, for states like WB and GJ, maximum percentage of sequences contains less than 5 co-mutations per sequence (Figure 3I-3J). Interestingly, in ‘Term3’ higher number of co-mutations are observed in sequences retrieved from both MH and TG (Figure 4D), maintaining the trend observed in ‘Term2’. For example, in TG more than 70% sequences harbored 8 and above mutations within single viral strain whereas in MH close to 45% sequences possessed 7 co-mutations. WB continued to harbor lower number of co-mutations even in ‘Term3’ with almost 60% sequences having 2 and 3 co-mutations per viral sequence.

**Figure 3:**
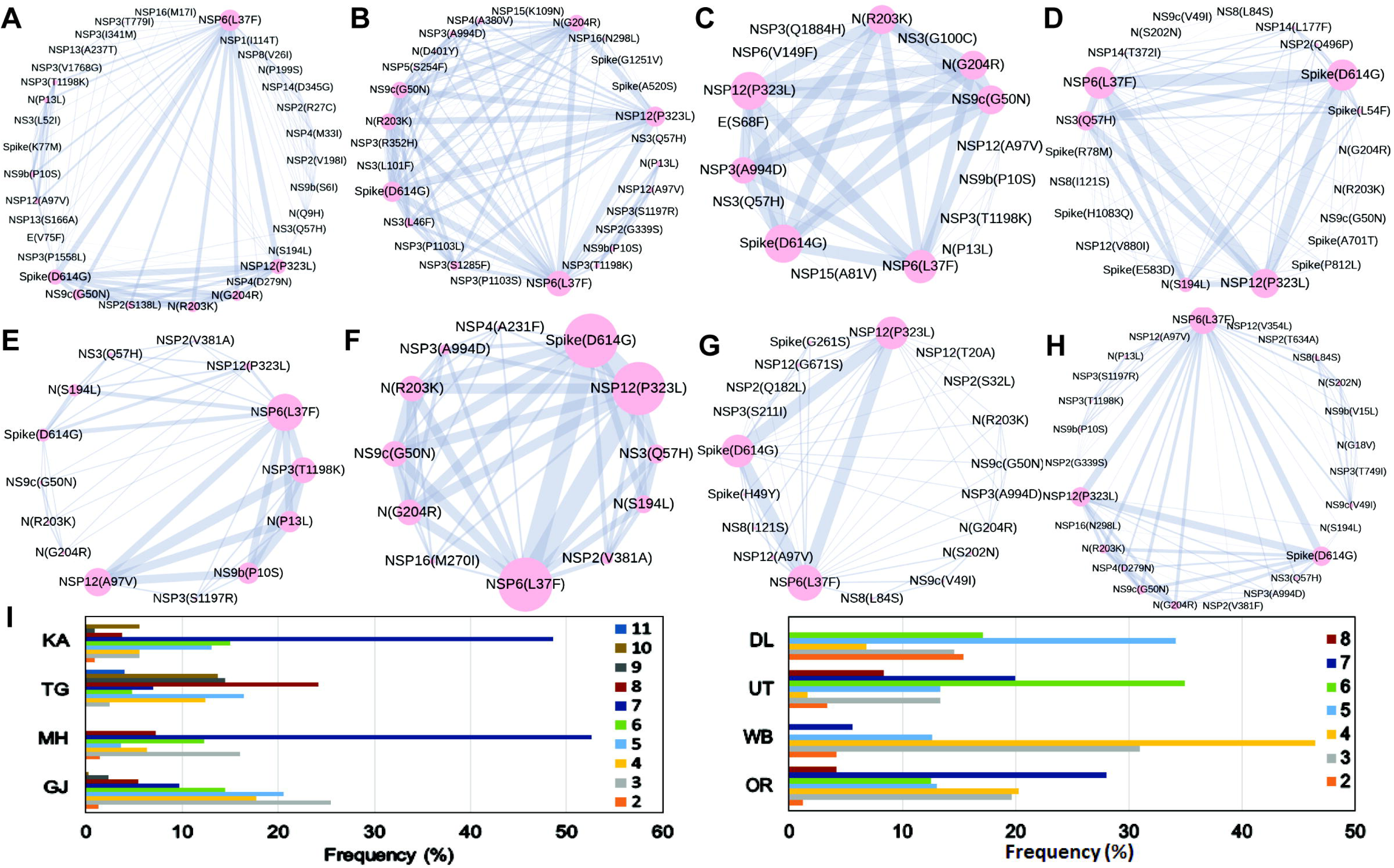
Co-mutation patterns and networks observed within Indian states and union territories during Term2. The networks of co-mutations for seven states and one UT, A. Karnataka (KA), B. Telangana (TG), C. Maharashtra (MH), D. Gujarat (GJ), E. Delhi (DL), F. Uttarakhand (UT), G. West Bengal (WB), H. Odisha (OR), where each mutation site is marked as a ‘node’ and the co-mutation is represented as an ‘edge’. Node size indicates the frequency of mutation while edge thickness represents the number of times (sequences) a pair of mutation is co-occurred. Panel I shows the bar plot distribution of the actual number of co-mutations within each state and UT.

**Figure 4:**
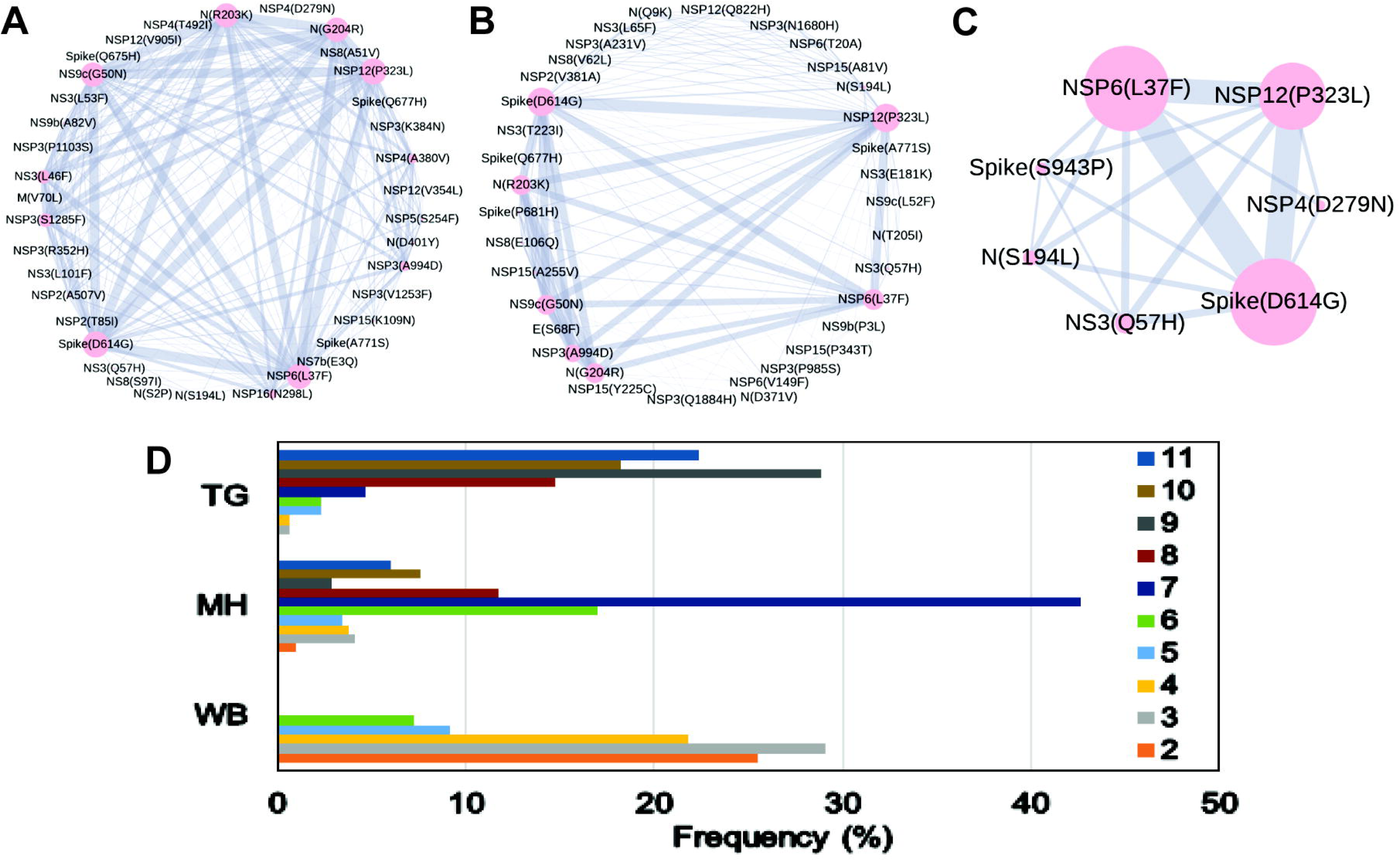
Co-mutation patterns and networks observed within Indian states and union territories during Term3. Panels A, B, and C show the networks of co-mutations for Telangana, Maharashtra, and West Bengal, respectively where each mutation site is marked as a ‘node’ and the co-mutation is represented as an ‘edge’. Node size indicates the frequency of mutation while edge thickness represents the number of times (sequences) a pair of mutation is co-occurred. Panel B shows the barplot distribution of the actual number of co-mutations within each state.

### Dynamics of co-mutation patterns in Indian states across ‘Term2’ and ‘Term3’

Similar to the comparison of frequent mutations across time scale, we decided to check whether the co-mutation patterns of SARS-CoV2 variant also varies over time within the states and union territories of India. However, as only three states possessed more than 50 sequences we could compare the changes in co-mutation patterns/motifs for these three states only along with overall Indian sequences. Panel A of Figure 5 compares the frequent mutations across three or two terms for sequences retrieved from India, Maharashtra (MH), Telangana (TG) and West Bengal (WB), respectively. As indicated before, significant fraction of novel mutations were evolved during ‘Term3’ compared to ‘Term2’ in India as well as in these three states. Comparison between the co-mutation networks consisting three to five (>=3 & <=5; Figure 5B upper panels) and more than five mutations (>5; Figure 5B lower panels) again shows emergence large number of unique co-mutation patterns and combinations for overall Indian sequences as well as sequences retrieved from MH, TG, and WB, respectively. This observation is further corroborated when we compared the co-mutation pattern within the major clades (G, GH, and GR, respectively) (Figure S5). GR clade has large number of co-occurred mutations of size (n) >5 in ‘Term3’ (Figure S5H) which is consistent with large number of co-occurred mutations of size (n) > 5 in MH and TG states (Figure 4D) where GR is the dominant clade (Figure S4A).

**Figure 5:**
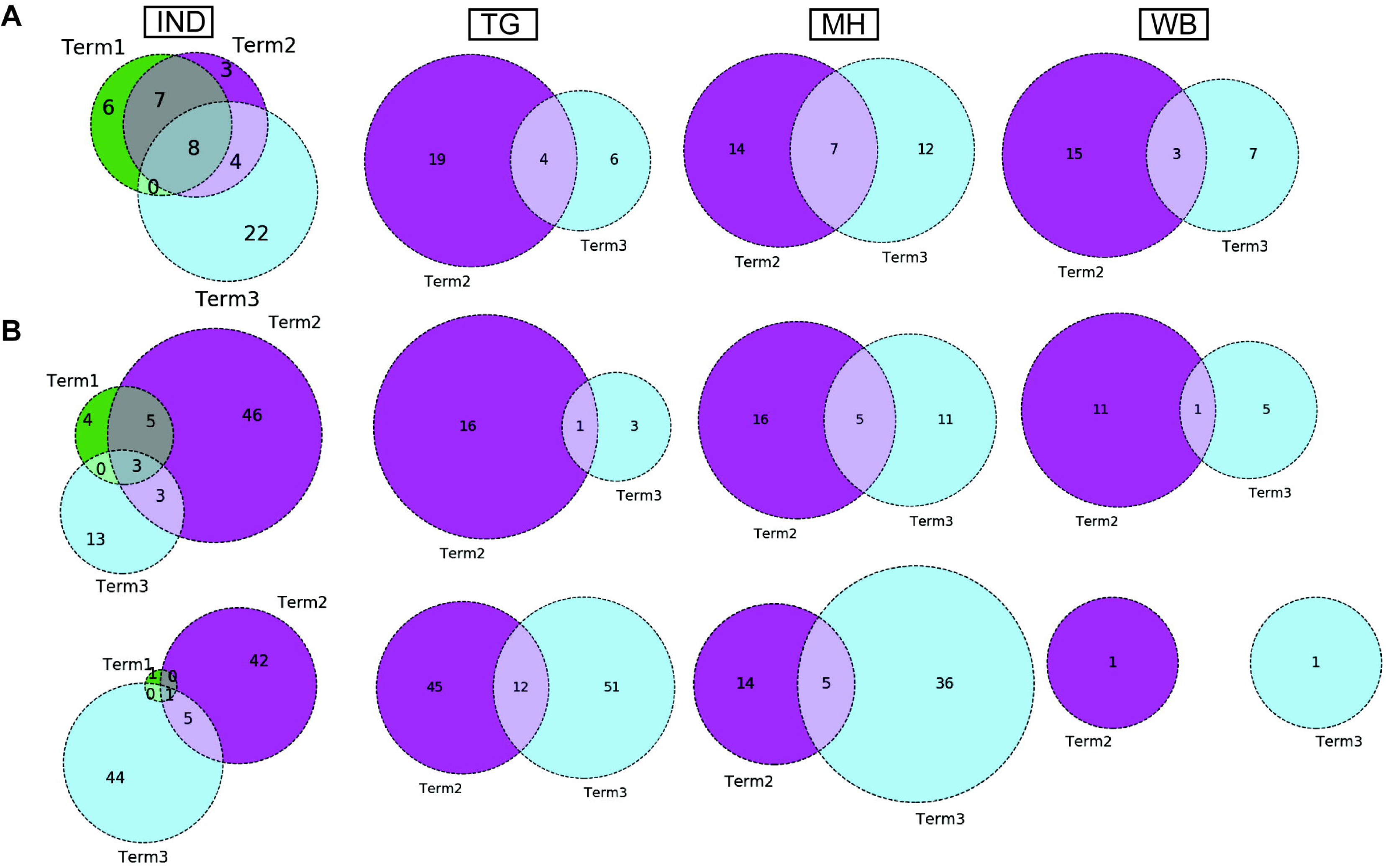
Dynamics of mutation and co-mutation pattern in Indian states across time spans. Panel A shows the overlap of frequent mutations observed in overall Indian samples (IND), Telangana (TG), Maharashtra (MH), and West Bengal (WB) collected across Term1 (green), Term2 (purple), and Term3 (cyan), respectively. Panel B: overlap of co-mutation patterns in three ‘Terms’ for where upper panel represents overlap of co-mutations having mutations between >=3 and <=5 sequences. Lower panel represents overlap for >=5 co-mutations per sequence.

### Association of specific mutations and co-mutations with patient status

Association of prevalent mutations with the status of the infected patients is a key aspect to explore the connection between genetic variability and the patho-physiology of the COVID-19 disease. Hence, we calculated the fraction of mutations that are predominantly found in a subset of Indian COVID-19 patients for which disease outcome information is available at the GISAID database. Out of the 3277 sequences deposited during 2020, patient status was reported for only 806 sequences where 95 were marked as deceased, and 631, 49, 31 were marked as symptomatic, mild, and asymptomatic, respectively. Figure 6 provides a matrix representation of the percentage of frequent mutations observed within the Indian patients marked as deceased (D), symptomatic (S), mild symptomatic (M), and asymptomatic (A), respectively. Statistical significance of the association of these mutations with respective categories was evaluated using Fisher’s Exact test (see Methods for details) and the list of mutations that are found to be significant (p-value ≤ 0.05) are listed in Table S2. It is evident that there is certainly some mutations [NS3(Q57H), N(S194L), Spike(L54F)] which are specifically associated with symptomatic and deceased status apart from the usual Spike(D614G) and NSP12(P323L) mutations (Figure 6). Among them NS3(Q57H), marker for GH clade is associated with deceased and symptomatic patients in North America also where GH is the dominant clade (Figure S2). NSP2(T85I) is also associated with deceased and symptomatic patients in North America but not in India.

**Figure 6:**
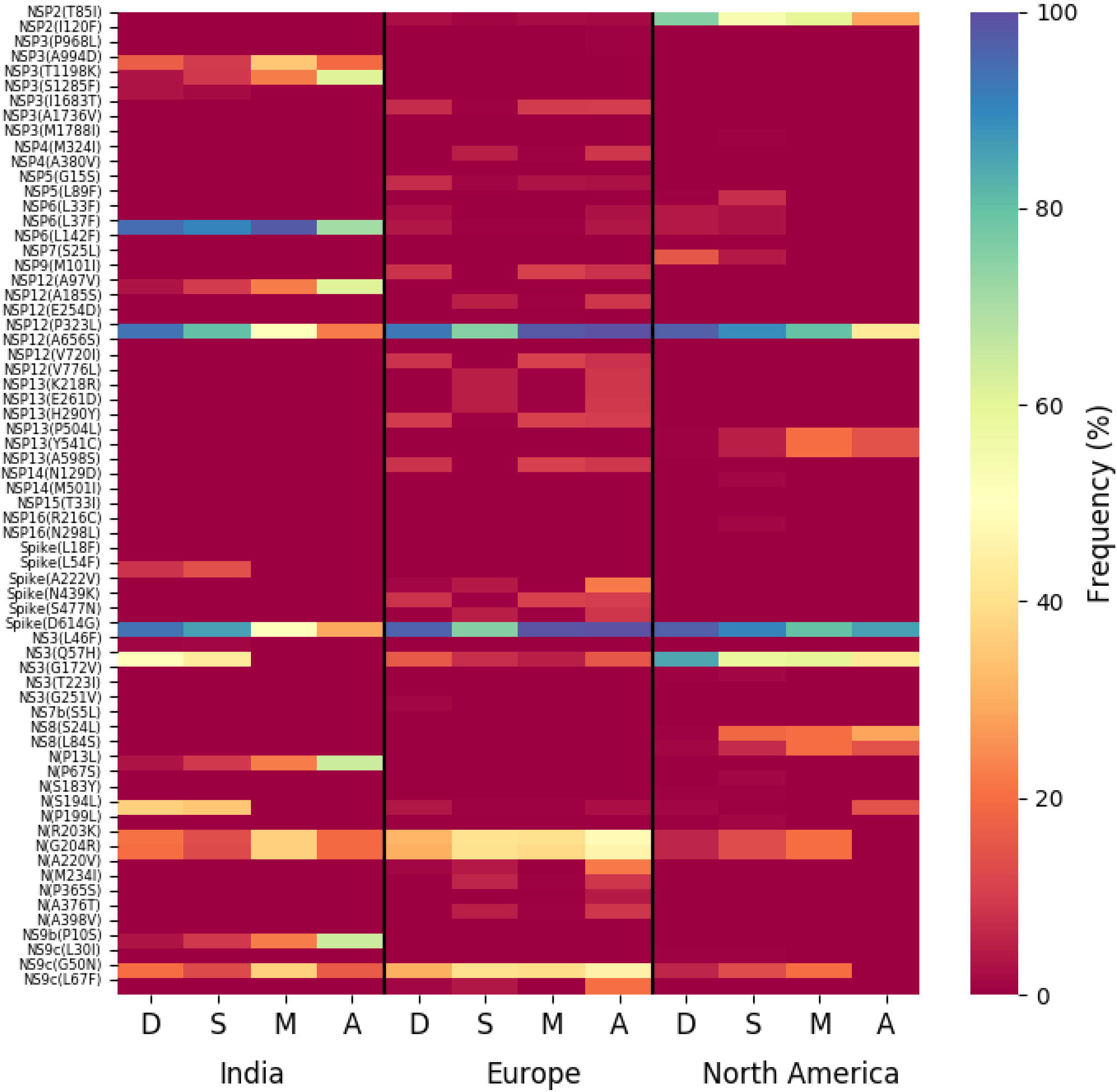
Association of specific mutations with the status of the COVID-19 patients from India, Europe and North America. A heat map of mutations and their frequencies that were found to be associated with four different categories of the COVID-19 patients’ status, Deceased (D), Symptomatic (S), Mild (M), and Asymptomatic (A), respectively are shown.

However, there are certainly some mutations that [NS9b(P10S), N(P13L), NSP12(A97V), and NSP3(T1198K), respectively) are relatively more specifically associated with asymptomatic patients in India. However, very few such asymptomatic specific mutations apart from N(S194L) was observed in North American frequent mutations (Figure 6). 15.54% of the deceased population and 3.91% of the symptomatic population of the N. American patients showed NSP7(S25L) mutation while missing in mild and asymptomatic populations. In Europe, a few frequent mutations like Spike(A222V), N(A220V), and NS9c(L67F) were observed relatively more within asymptomatic patients (Figure 6). Examination of association of clade specific frequent mutations with the disease/patient status does not reveal much specific association. However, relatively higher (>=10%) association was observed for a G clade (Indian population) mutation, N(S194L), which was exclusively found in 16% symptomatic patients only (Figure S6A). In case of GH clade, N(S194L) is observed in more than 70% samples of deceased and symptomatic patients. Similarly, NSP14(T372I), a GH clade specific mutation (Indian population) exclusively observed in symptomatic patients with higher frequency (10%) but not in deceased population. NSP2(T85I) does not show any specific association with any patient status in overall European patients (Figure 6), however within GH clade, it is reported in 50% mild patients (Figure S6B). European GR clade specific NSP3(K945N), Spike(L5F), Spike(S549Y), Spike(M1229I), and NS9b(R13L) mutations show more than 10% abundance in mild patients only (Figure S6B). NSP13(S485L), NSP14(T250I) and NS3(S74F) from GH clade, NSP3(P340S), NSP3(I414V), NSP13(A505V), NS3(W131C), N(D377Y) from GR clade, and NSP3(T1456I), NSP4(T492I), NSP14(T113I), Spike(A262S), Spike(P272L), Spike(G639S), NS7a(Q94L), NS8(I121L), N(P365S) from GV clades seem to be associated with asymptomatic patients only (Figure S6B). In N. American GH clade specific frequent mutation NSP14(A320V) was found exclusively only in deceased (16%) whereas NSP5(L89F) and NS8(S24L) were observed in 14% and 32% symptomatic patients, respectively (Figure S6C).

Impact of the single point mutation could be essential but may not be sufficient to elicit pathological response and or variation in disease phenotype. Hence, we thought of exploring the association of evolutionary selected multiple mutations within a viral strain with respect to the disease status of the patient from whom the strain was isolated. Figure 7 presents the frequency of patients infected with SARS-CoV2 strains that harbors the co-mutation combinations. Combination of five co-mutations [NSP3(T1198K)-NSP6(L37F)-NSP12(A97V)-NS9b(P10S)] were found to be significantly higher (>50%) in asymptomatic patients whereas co-mutations of NSP3(A994D)-NSP6(L37F)-NSP12(P323L)-Spike(D614G)-N(G204R)-N(R203K)-NS9c(G50N) and NSP6(L37F)-NSP12(P323L)-Spike(D614G)-NS3(Q57H)-N(S194L) were observed in higher frequencies in mild patients (Figure 7). However, it is evident that co-mutations of NSP6(L37F)-NSP12(P323L)-Spike(D614G) is present as either triad or part of larger co-mutation network in most of the symptomatic patients along with mild, symptomatic and deceased populations. However, quite different co-mutation patterns are observed with patients from Europe and N. America where absence of NSP6(L37F) mutation was along with Spike(D614G) and NSP12(P323L) mutations (Figure S7).

**Figure 7:**
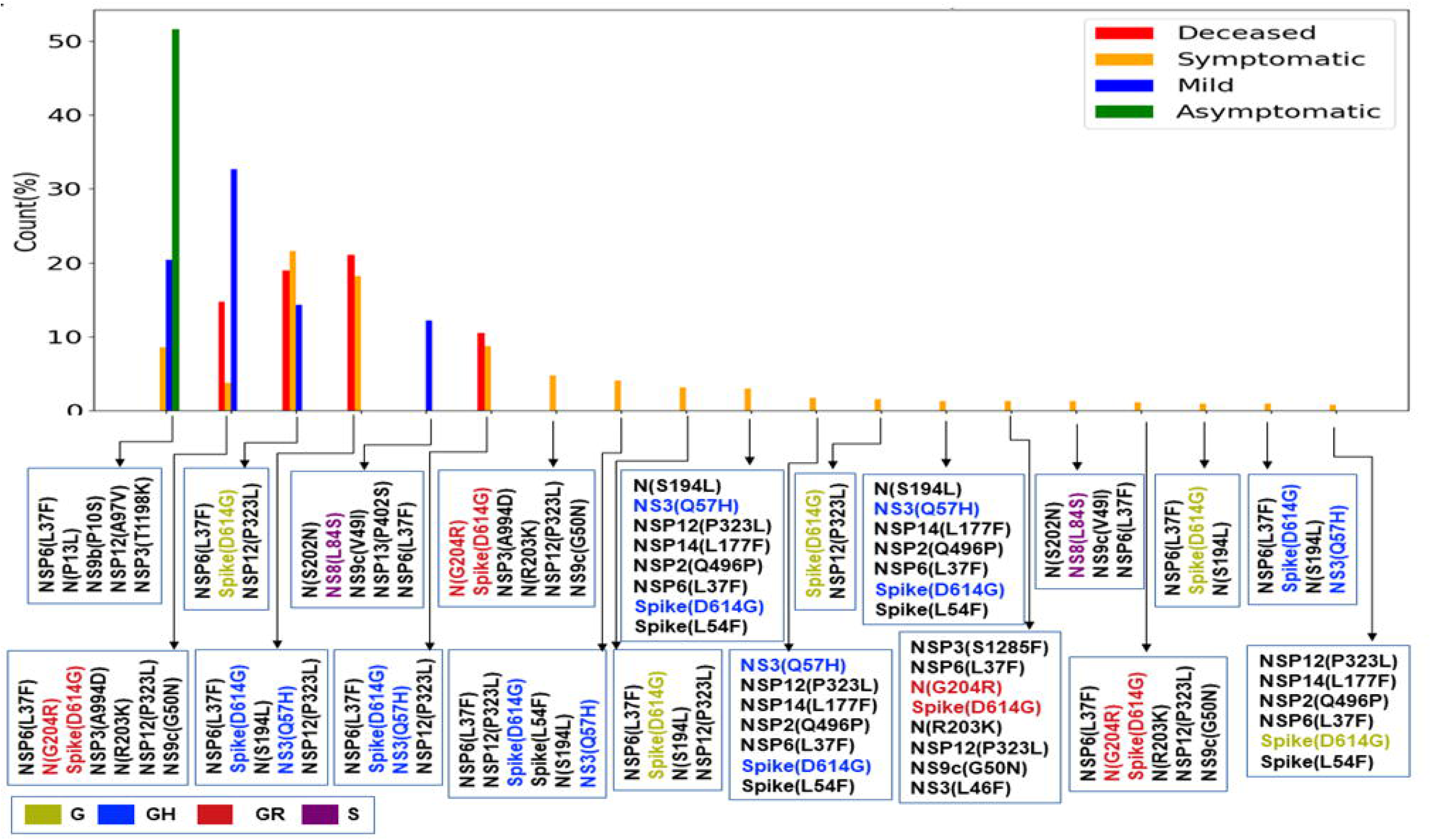
Association of specific co-mutation patterns with the status of the COVID-19 patients from India. Frequencies of specific co-mutations are plotted with respect to the four COVID-19 patients’ status categories Deceased, Symptomatic, Mild, and Asymptomatic, respectively. G, GH, GR and S clades defining mutations are marked in yellow, blue, red, and purple, respectively.

## Discussion

Whole-genome sequencing of SARS-CoV2 strains throughout the world has provided enormous amount of knowledge about the evolutionary diversity of this deadly virus and also contributed significantly in understanding the nature of the pandemic (35, 36). During the difficult times in late 2019 and throughout 2020, sequencing of the viral strains was one of the primary research objectives globally in order to understand the specific phylogenetic variations and their connection to the spread and transmissibility of the virus (37). As part of this global endeavor, deposition of SARS-CoV2 genome data in genomic databases such as GenBank (38), NCBI (39), and GISAID (16) has truly facilitate the knowledge to tackle the deadly COVID-19 disease. Here, 54 SARS-CoV2 whole genomes have been sequenced using NGS platforms with very high coverage and depth (Figure S1B). The viral RNA was extracted from patients from the city of Kolkata, capital of West Bengal (WB) state of India. These 54 RNAs were collected during the time period of August-October 2020 and constitute almost all the SARS-CoV2 (except one) genome sequences deposited from West Bengal (WB) in the latter half of the year 2020. However, the clade distributions of these 54 sequences constituting the ‘Term3’ repertoire of WB sequences are distinctively different than that of overall India (Figure S1 and Figure 1). WB ‘Term3’ sequences have significant higher fractions of ‘O’ clade (GISAID) and 20A clade (Nextstrain) sequences compared to both overall Indian distribution and sequences retrieved during the ‘Term3’ period as well. As a matter of fact the ‘Term2’ (April 2020 – July 2020) sequences from WB show significantly higher fraction of ‘G’ and 20A clades distribution (82% and 83%, respectively) compared to other states and overall India (Figure 1 and 2). This indicates that the evolutionary dynamics of the WB strains are slower and therefore may be the usual fixation of major clades like ‘GR’ or 20B is delayed for some reason.

Intrigued by this observation, we thought it would be helpful to undertake an analysis to examine the variations in evolutionary dynamics of the SARS-CoV2 virus and its connection to the spatio-temporal regulation. In other words, we thought it would be worthwhile to check the evolutionary changes in viral populations prevalent in India across time and geographical zones. Our analysis clearly shows gradual fixation of the more prevalent clades such as GR (GISAID) and 20B (Nextstrain) across time within the viral population extracted from Indian patients. However, the most frequent mutations observed within the population across three time terms were quite different and emergence of newer mutations were observed even at the end of the year 2020 for the overall Indian population as well as population categorized based on their clades. Similarly, sequences belonging to same clades but extracted from different states also show little commonality in the type of non-clade defining mutations (Figure 2, 3 and Table 2).

Further, we also examined the variation in co-occurrence of mutations (co-mutation) across Indian states and union territories compared over the three time terms. We observed a large number of non-clade defining mutations to co-occur within SARS-CoV2 strains from the states and union territories during the ‘Term2’ and ‘Term3’ which is consistent with earlier reports based on the sequences collected mostly in ‘Term1’ period (17). Consequently, a larger number of unique combinations among these co-occurred mutations were appeared during the latter half of the year 2020. It is really interesting to observe that larger fractions of viral strains extracted from states like Telangana, Maharashtra and Karnataka during ‘Term2’ harbored five or more co-mutations while major fractions from West Bengal and Gujarat possessed less than five co-mutations. Even in ‘Term3’ far larger fractions of sequences from Telangana and Maharashtra showed higher co-mutating residues compared to that of West Bengal. Significantly higher numbers of active infection cases were reported in Maharashtra and Karnataka compared to West Bengal and Gujarat during ‘Term2’ and ‘Term3’ (15, 40). Hence, the higher number co-mutations in those two states could be associated with higher infectivity of the strains prevalent there. In West Bengal during ‘Term3’ only ∼22% sequences harbored more than 5 co-mutations while almost 58% of the sequences were designated as ‘O’ clade known for harboring larger amount of other (34) mutations.

Finally, we tried to associate the frequent mutations and co-mutation patterns along with COVID-19 patient status broadly categorized into deceased, symptomatic, mild, and asymptomatic groups, respectively. Although the number of patient status mapped data is lower (806 out of 3277 sequences) we thought it was worthwhile to investigate if some of the mutations and co-mutation patterns could be specifically associated with those four states of the COVID-19 disease. Apart from the usual Spike(D614G) and NSP12(P323L) mutations, specific association of mutations NS3(Q57H), N(S194L), and Spike(L54F) are observed with symptomatic and deceased status of Indian patients while NS9b(P10S), N(P13L), NSP12(A97V), and NSP3(T1198K), respectively are found to be relatively more in asymptomatic Indian patients. Similarly, presence and absence of specific mutation with respect to symptomatic or asymptomatic status were also examined for North American and European samples for which the disease status was available at the GISAID database. Although we did not find any specific co-mutation pattern with severe (deceased or symptomatic) patients but few co-mutation patterns were observed to be significantly higher in asymptomatic and mild symptomatic patients (Figure 7). We have also identified presence of a co-mutation triad [NSP6(L37F)-NSP12(P323L)-Spike(D614G)] in most of the symptomatic patients including mild, symptomatic and deceased populations of Indian patients. Figure 8 summarizes the most frequent mutations and co-mutation network motifs within overall Indian SARS-CoV2 samples, most prevalent clades and also for the viral strains extracted from patients for which the disease status was documented. We observed the top 5 mutations for deceased and symptomatic patients remain identical. Among these 5 mutations, 3 appears in mild and only one mutation (NSP6(L37F)) appears in asymptomatic patients. It indicates the possibility of association of specific mutations with disease severity. Similar overlap is observed in the co-mutation networks of deceased and symptomatic patients. The co-mutation network observed in 51% of asymptomatic patients is also observed in 20.41% mild patients but neither in deceased nor in case of symptomatic patients. Similarly, Figure 9 provides an overview of the dynamics of co-mutation pattern/motif observed within viral strains from Indian population, most prevalent clades and for three states (MH, TG and WB) for which sufficient sequences were deposited in GISAID database in Term2 and Term3. It highlights emergence of new co-mutation network motifs with time across different states and clades. We believe that our report is one of the few studies that aim to provide a comprehensive picture of the evolutionary dynamics and co-mutations patterns of the SARS-Cov2 strains prevalent within Indian population throughout the year 2020.

**Figure 8:**
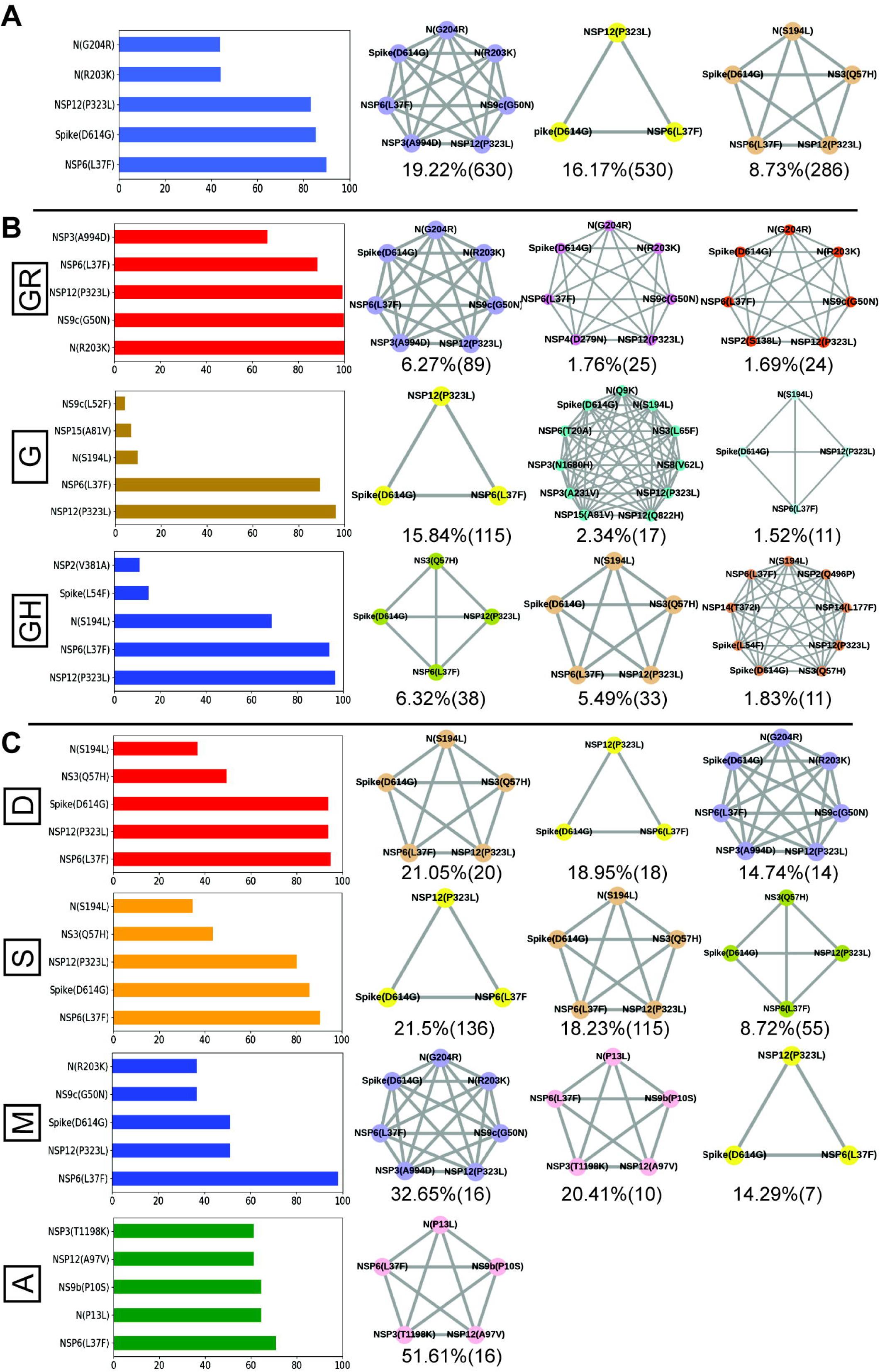
Frequent mutations, co-mutations and their association with disease severity in India. A. Most frequent 5 mutations and 3 co-mutation patterns in India. B. Most frequent 5 mutations and 3 co-mutations in major clades of India. Clade defining mutations are excluded in the bar plot. C. Most frequent 5 mutations and 3 co-mutations in different types of patients. Different color codes are used for specific co-mutation network motifs. Network motifs were sorted and ranked exclusively.

**Figure 9:**
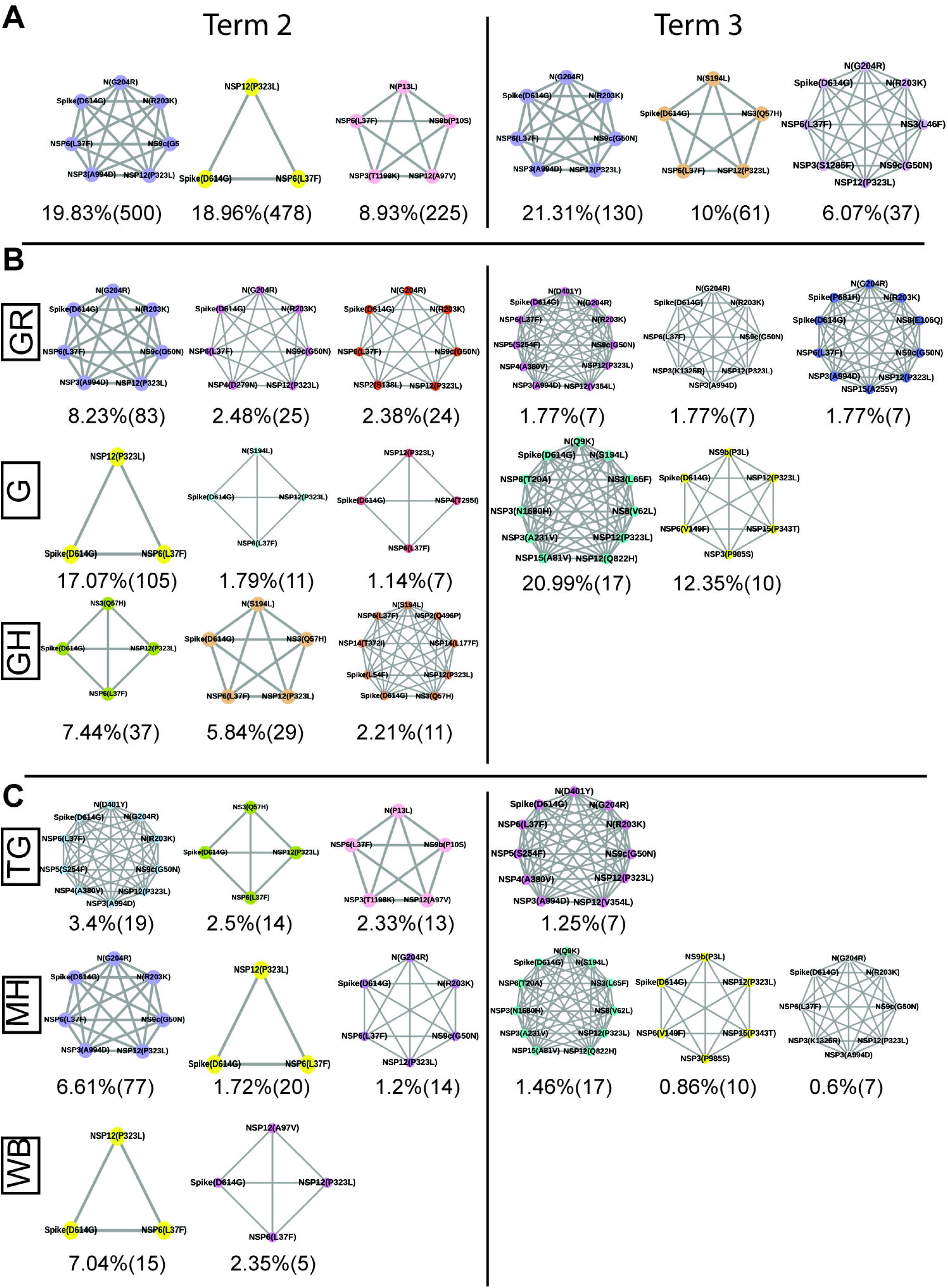
Spatio-temporal variation in most frequent co-mutations in India. A. Most frequent 3 co-mutations in Term2 and Term3 in India. B. Most frequent 3 co-mutations in Term2 and Term3 in major clades of India. C. Most frequent 3 co-mutations in Term2 and Term3 in different states of India. Co-mutation network observed in at least 5 patients are represented. Different color codes are used for specific co-mutation network motifs. Network motifs were sorted and ranked exclusively.

## Authors’ contributions

NB performed all the sequence analysis, mutational and co-mutational analysis. PM and SKM sequenced the viral genomes. DB assembled the raw sequences and performed computational analysis. SS, AR, and AGM provided clinical guidance and performed RNA processing and detection of COVID-19 from bio specimens. SC, SP and PC conceptualized and coordinated the project. NB and SC wrote the manuscript.

## Acknowledgements

The authors acknowledge CSIR-Indian Institute of Chemical Biology for infrastructural support. SC acknowledges financial support from CSIR MLP-132 grant. NB acknowledges the Systems Medicine Cluster (SyMeC) grant (GAP357), Department of Biotechnology (DBT), Government of India for fellowship. PM acknowledges University Grants Commission, Government of India for her fellowship. DB thanks Indian Council of Medical Research, New Delhi for a senior research fellowship. SP was supported by Ramanujan Fellowship of Science and Engineering Research Board (SERB), Government of India. The funders had no role in study design, data collection and analysis, decision to publish, or preparation of the manuscript.

## Supplementary Figure legends

**Figure S1:** Q**u**ality**, demographic and phylogenic analysis of the SARS-CoV2 genome sequences.** Panel A compares the age distribution of the patients from which the 54 SARS-CoV2 sequenced strains (IICB) were extracted with respect to the overall Indian COVID-19 patients’ (India) age distribution.

Panel B show the depth (shown as multiplies of ‘X’) of the sequenced data for both long (Nanopore) and short (Illumina) read sequences.

Panels C and D plot the GISAID (left) and Nextstrain (right) clade distribution of the 54 genomes sequenced by CSIR-IICB and MEDICA Superspecialty Hospital and sequenced deposited from overall India, respectively.

**Figure S2: Distribution and dynamics of SARS-CoV2 clades in the world for three different time spans in the year 2020.** Panels A, B, and C show distribution of GISAID clades across the world for three different times spans, ‘Term1’ (December 2019 to March 2020), ‘Term2’ (April 2020 to July 2020) and ‘Term3’ (August 2020 to December 2020), respectively.

**Figure S3: Overlap of frequent mutations across three terms for most prevalent GISAID and Nextstrain clades in India.** Panel A shows the overlap of frequent mutations observed in the most prevalent GISAID clades like G, GH and GR collected across Term1 (green), Term2 (purple), and Term3 (cyan), respectively.

Panel B shows the overlap of frequent mutations observed in the most prevalent Nextstrian clades like 19A, 20A and 20B collected across Term1 (green), Term2 (purple), and Term3 (cyan), respectively.

**Figure S4: Distribution of GISAID clades within Indian states and comparison of frequent mutations across them during Term3 (August 2020 to December 2020).** Panel A shows the distribution of GISAID clades in three Indian states that deposited more than 50 SARS-CoV2 sequences during August 2020 to December 2020.

Panels B-D plot the number of sequences and frequent mutations of the three states and show the number of common mutations among them for G, GH, and GH clades, respectively.

**Figure S5: Dynamics of co-mutation pattern across three terms for most prevalent GISAID clades in India.** Overlap of co-mutation patterns in three ‘Terms’ for co-mutations having mutations between >=3 and <=5 sequences (panel A) and for >=5 co-mutations per sequence (panel B) are shown.

**Figure S6:** Association of specific mutations with the status of the COVID-19 patients from India, Europe and North America infected with specific clades of the **SRAS-CoV2 virus.** A heat map of mutations and their frequencies that were found to be associated with four different categories of the COVID-19 patients’ status, Deceased (D), Symptomatic (S), Mild (M), and Asymptomatic (A), respectively are shown for most prevalent clades.

**Figure S7: Association of specific co-mutation patterns with the status of the COVID-19 patients from Europe and North America.** Frequencies of specific co-mutations are plotted with respect to the four COVID-19 patients’ status categories Deceased, Symptomatic, Mild, and Asymptomatic, respectively. Panel A shows data from Europe while panel B is for North America. Clade defining mutations are marked in corresponding colors.

