## Supplementary figures and images for "Genomic surveillance and phylodynamic analyses reveal emergence of novel mutation and co-mutation patterns within SARS-CoV2 variants prevalent in India"

### Supplemental Figure 1

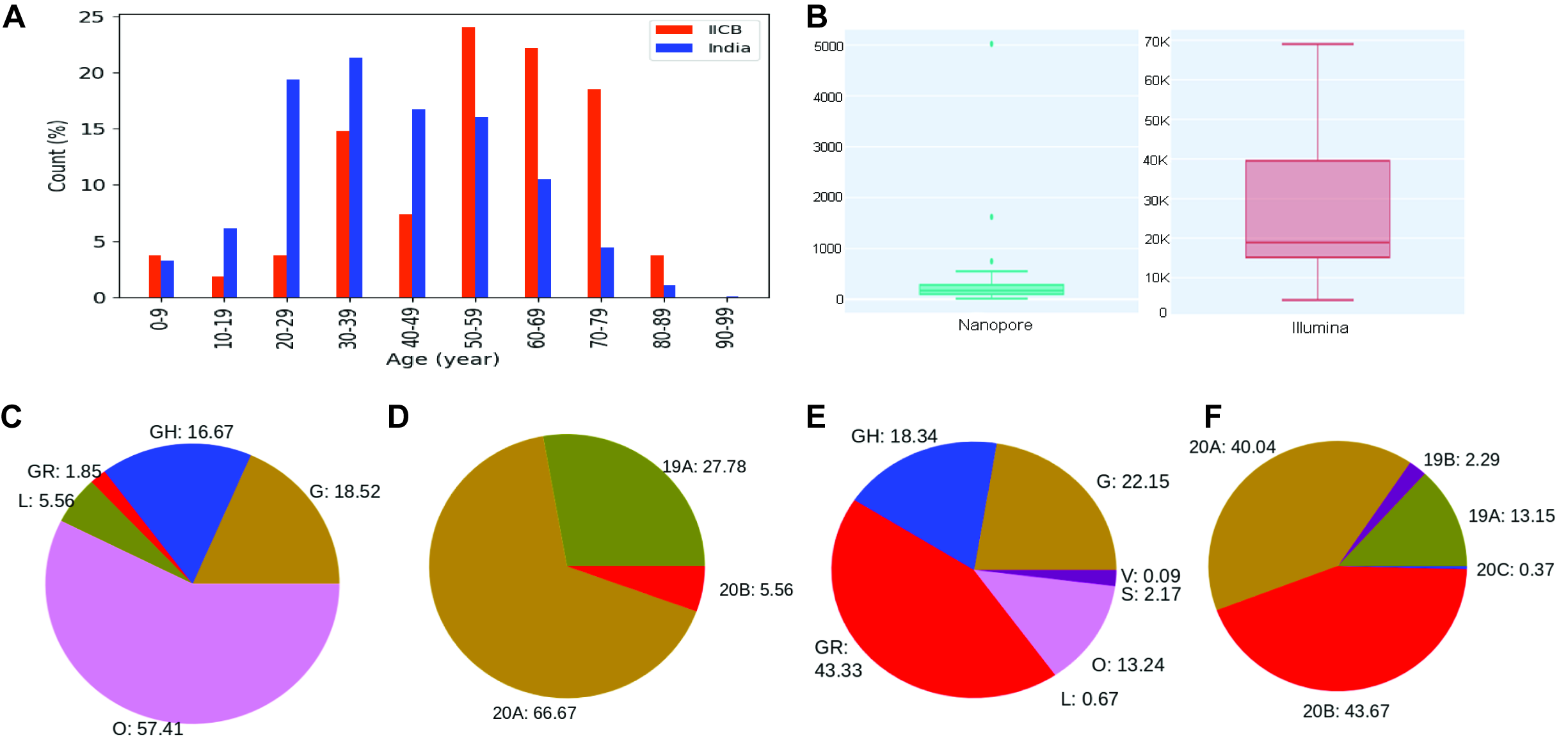

### Supplemental Figure 2

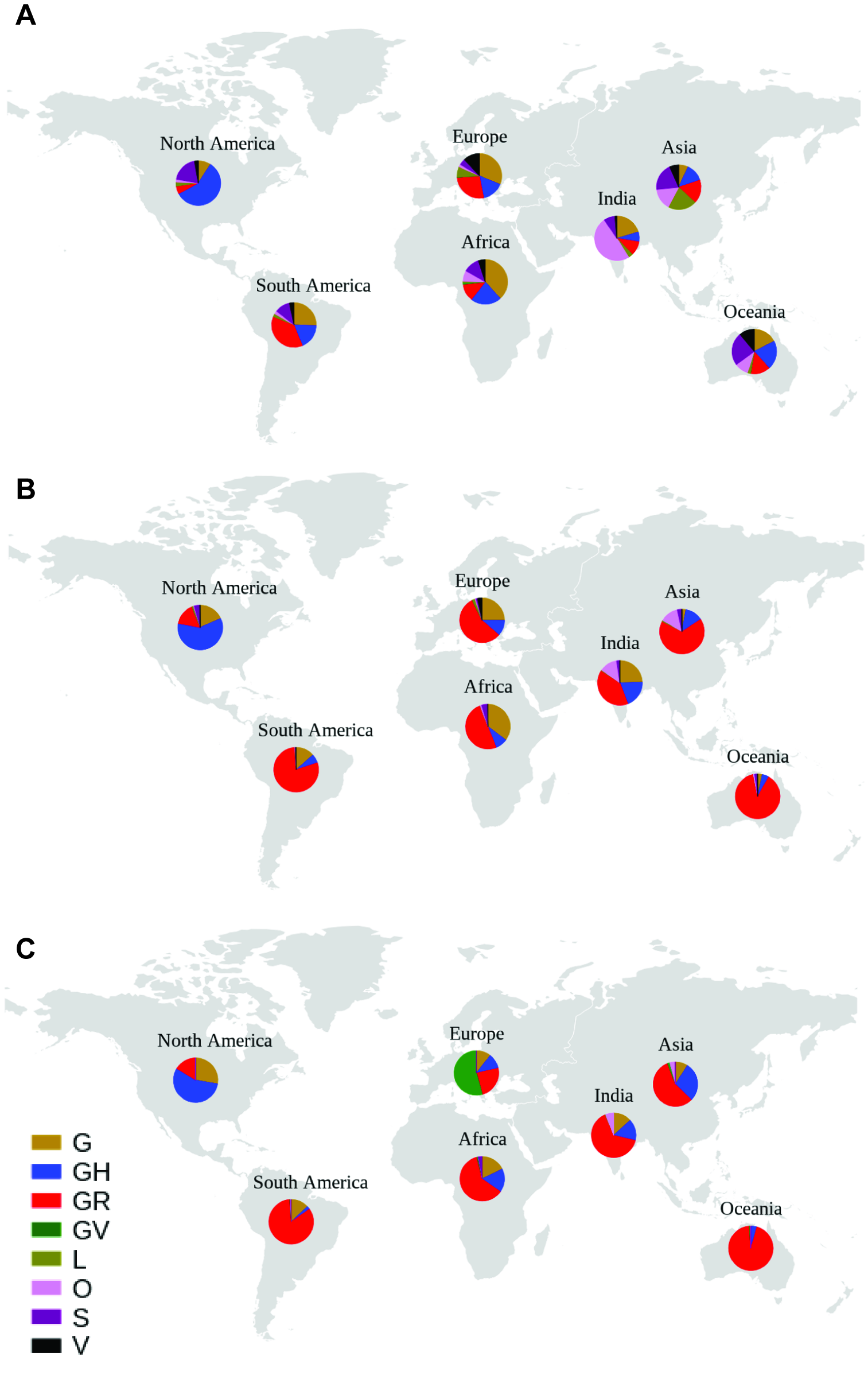

### Supplemental Figure 3

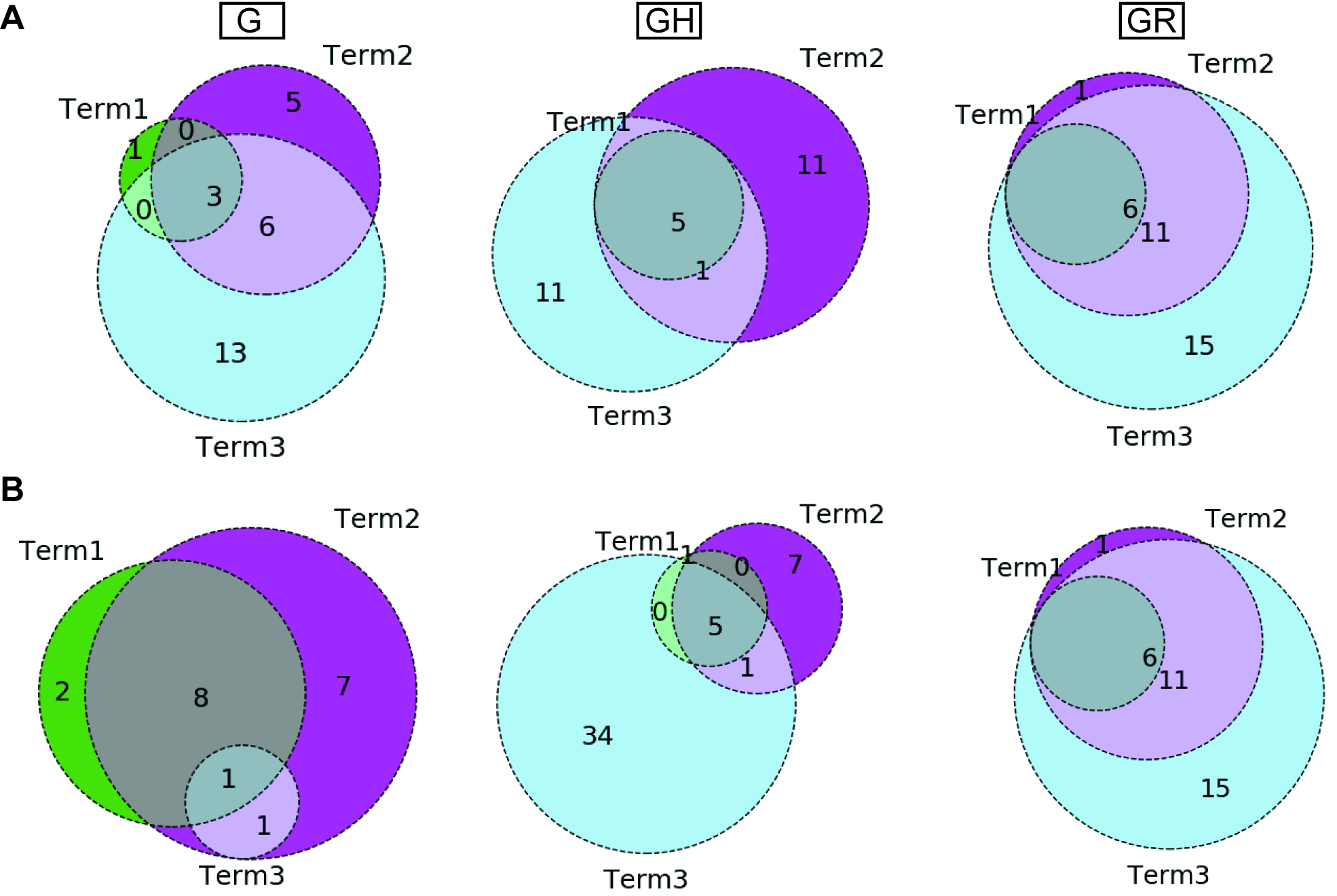

### Supplemental Figure 4

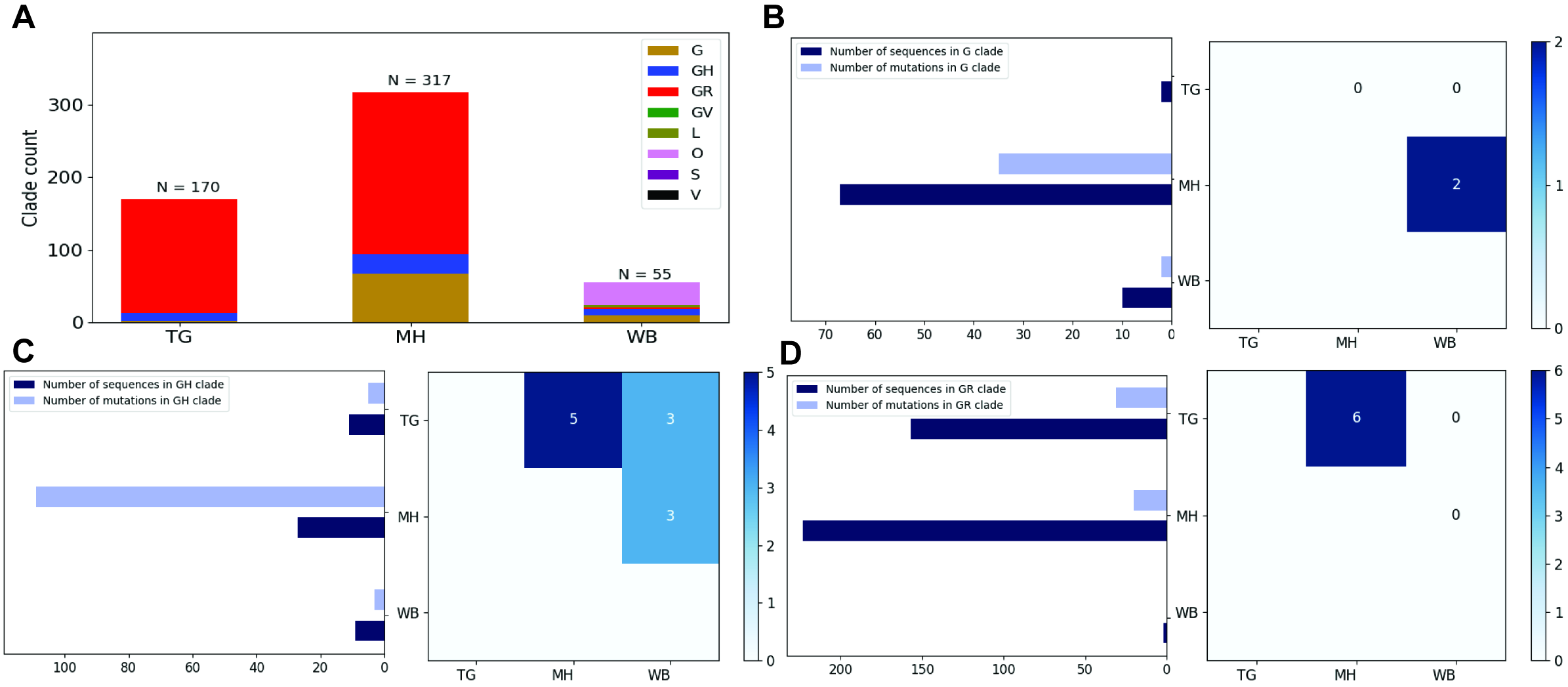

### Supplemental Figure 5

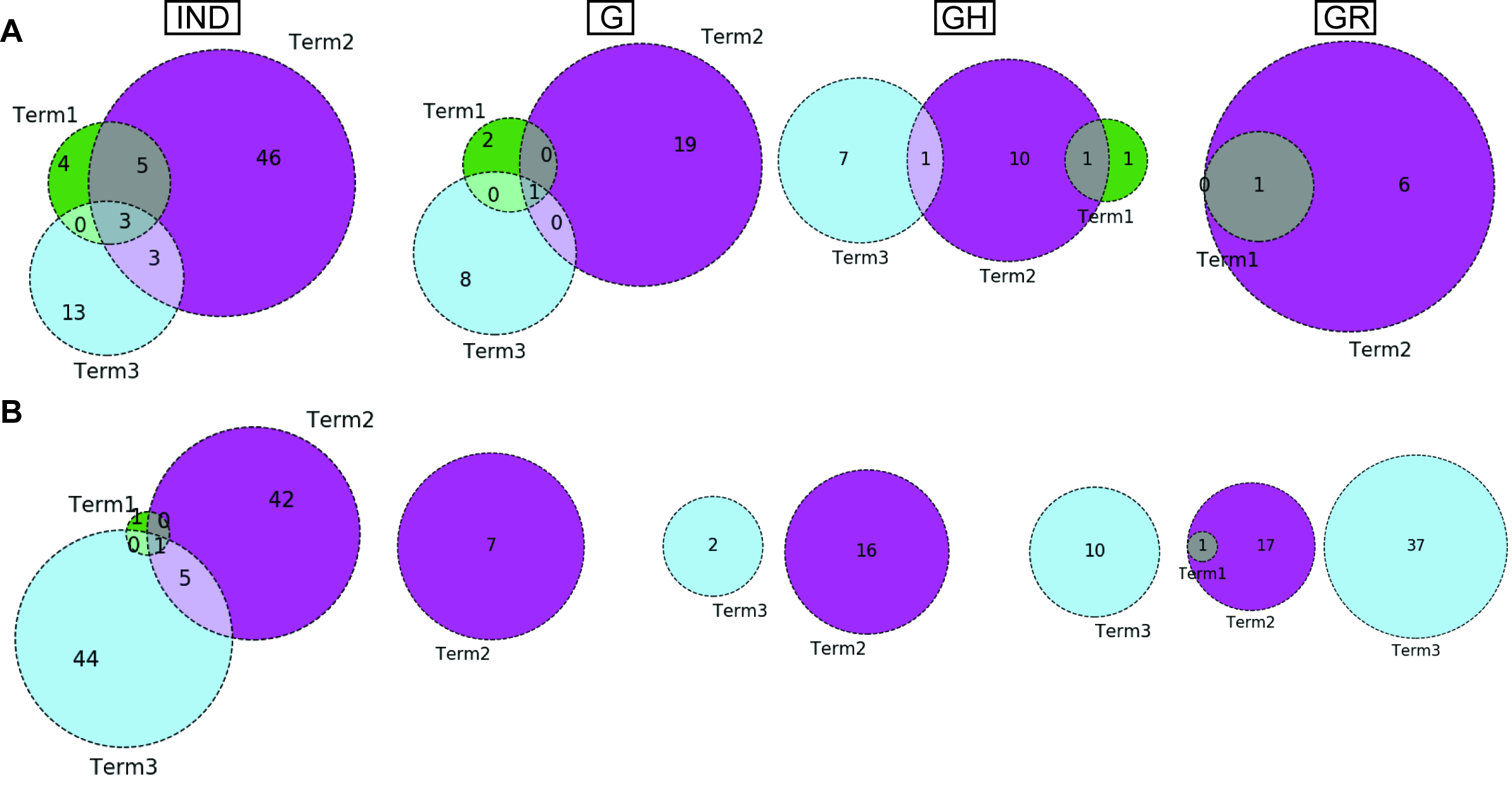

### Supplemental Figure 6

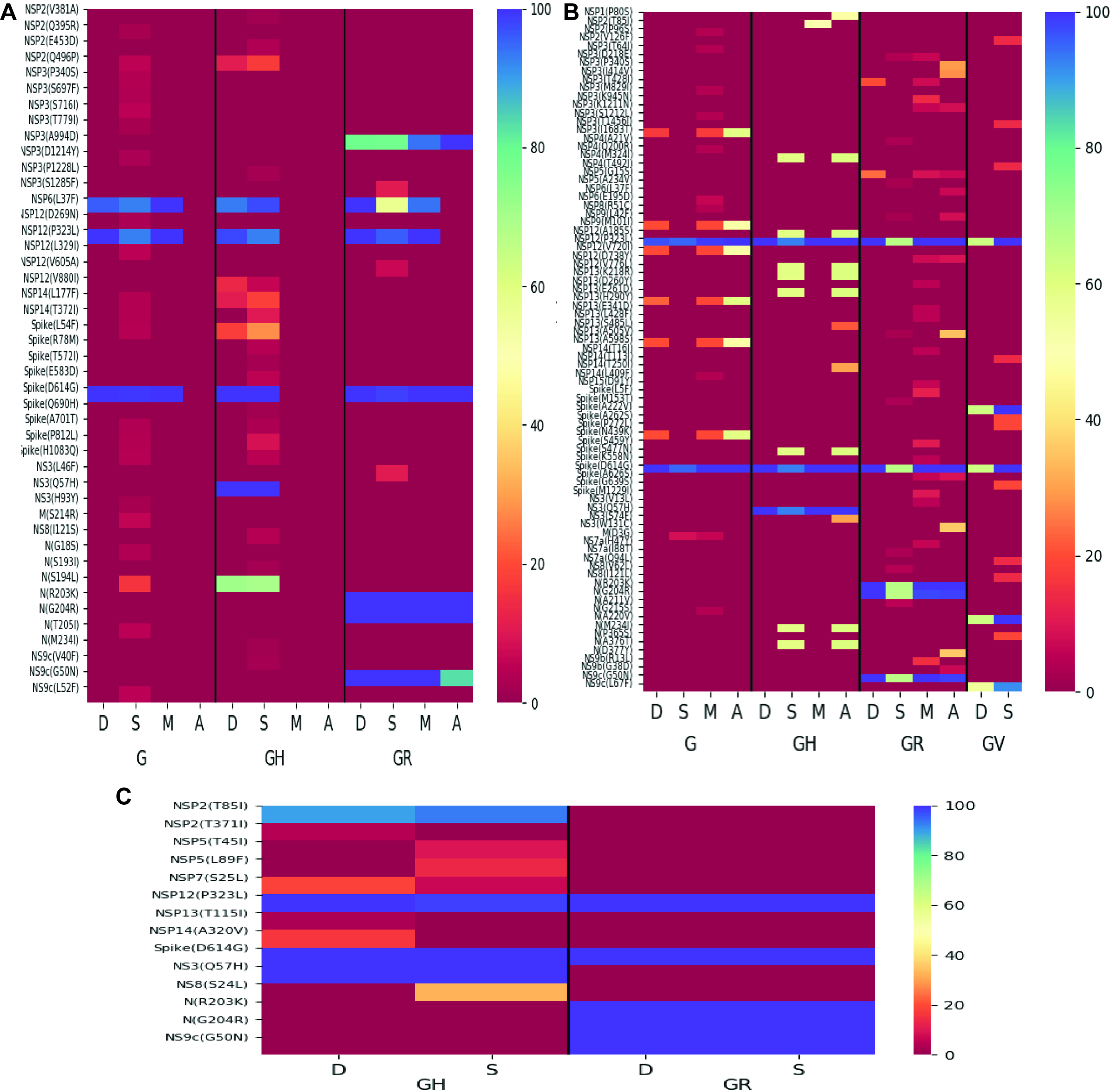

### Supplemental Figure 7

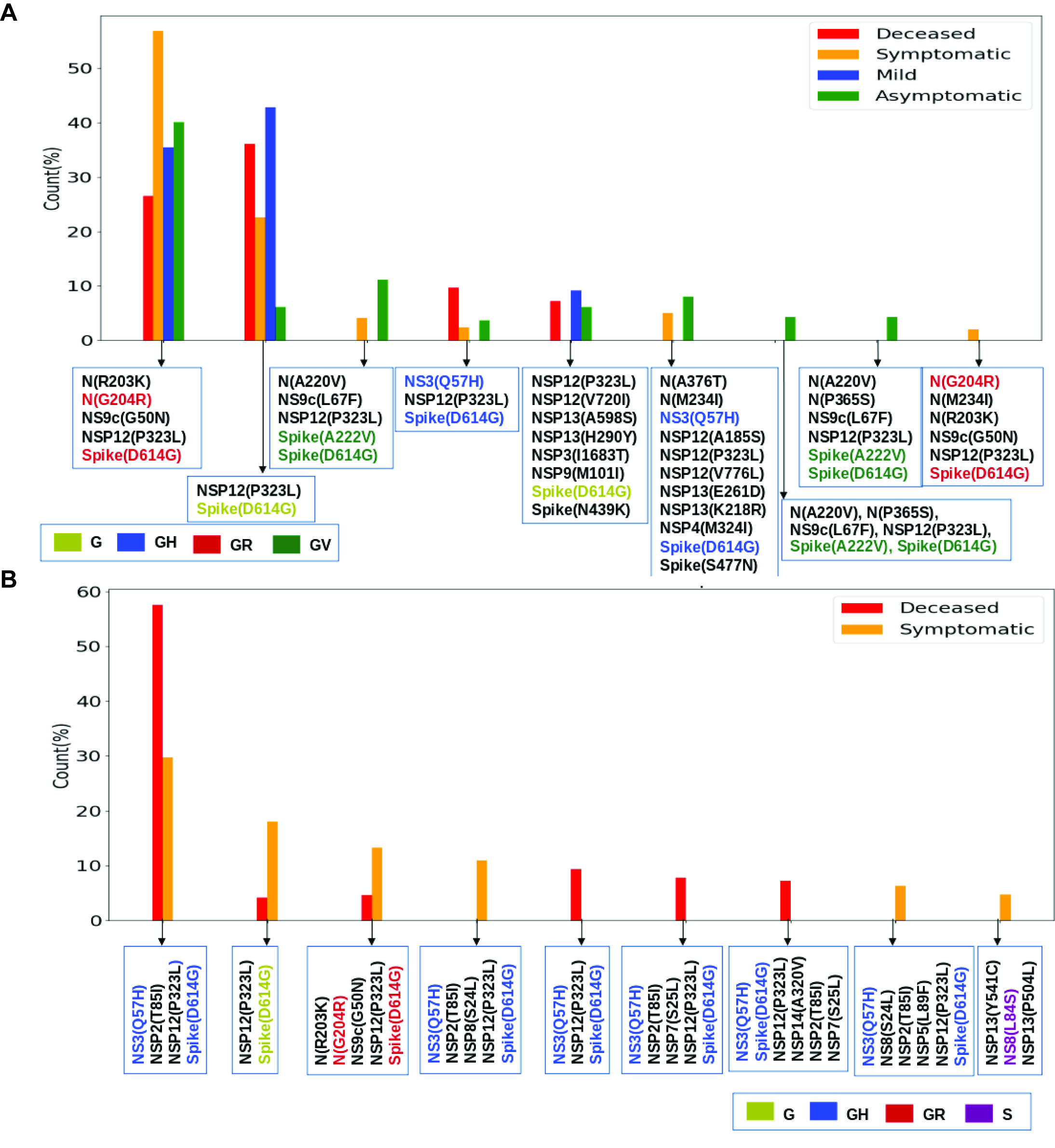
